# Adolescent ethanol exposure decreases cholinergic markers, blunts behaviorally-evoked acetylcholine and increases in apical dendritic branching within the orbital frontal cortex

**DOI:** 10.1101/2021.03.26.436863

**Authors:** BT Kipp, PT Nunes, E Galaj, B Hitchcock, T Nasra, KR Poynor, SK Heide, NL Reitz, LM Savage

## Abstract

During adolescence, heavy binge-like ethanol consumption can lead to frontocortical structural and functional impairments. These impairments are likely driven by adolescence being a critical time point for maturation of brain regions associated with higher-order cognitive functioning. Rodent models of heavy binge-like ethanol exposure show consistent disruptions to the typical development of the prefrontal cortex (PFC). All deep cortical layers receive cholinergic projections that originate from the Nucleus basalis of Meynert (NbM) complex. These cholinergic projections are highly involved in learning, memory, and attention. Adolescent intermittent ethanol exposure (AIE) induces cholinergic dysfunction as a result of an epigenetic suppression of the genes that drive the cholinergic phenotype. The current study used a model of AIE to assess structural and functional changes to the frontal cortex and NbM following binge-like ethanol exposure in adolescence. Western blot analysis revealed long-term disruptions of the cholinergic circuit following AIE: choline acetyltransferase (ChAT) was suppressed in the NbM and vesicular acetylcholine transporter (VAChT) was suppressed in the orbitofrontal cortex (OFC). In vivo microdialysis for acetylcholine efflux during a spatial memory task determined changes in cholinergic modulation within the PFC following AIE. However, AIE spared performance on the spatial memory task and on an operant reversal task. In a second study, Golgi-Cox staining determined that AIE increased apical dendritic complexity in the OFC, with sex influencing whether the increase in branching occurred near or away from the soma. Spine density or maturity was not affected, likely compensating for a disruption in neurotransmitter function following AIE.

**Significance Statement:** Adolescent ethanol exposure decreases cholinergic markers in the basal forebrain to orbital frontal cortical circuit, which lead to a massive suppression of behaviorally-activated tonic release acetylcholine and sex-dependent increases in dendritic branching. We concluded that cholinergic dysfunction is a contributor to cognitive impairments associated with heavy alcohol exposure during adolescence.

**Data Sharing:** The data that support the findings of this study are available from the corresponding author upon reasonable request.

## Introduction

Heavy binge-like alcohol exposure during adolescence, in humans and animal models, can manifest in frontocortical structural and functional impairments.^1–4^ Such disruptions are likely driven by the maturation of brain regions associated with higher-order cognitive functions that occurs in adolescence.^5–8^ Subsequently, it is not surprising that heavy binge drinking during adolescence has been associated with impairments in executive functions.^9,10^ In rodent models, heavy binge-like ethanol exposure impacts the typical development of the prefrontal cortex (PFC). There are persistent suppressions of PFC cortical volume and myelination, whereas markers of neuroinflammation and neurodegeneration are increased.^11–15^ Such models also revealed that heavy binge-type ethanol exposure produces deficits in set-shifting^16,17,^ working memory^18,19,^ behavioral inhibition^20,^ and reversal learning.^12,13,21^ These aberrant behaviors are classified under the domains of cognitive and behavioral flexibility and result in an inability to readily adapt to changes in the environment^22,23^

A key neural alteration that occurs after heavy alcohol exposure during adolescence, is the disruption of the forebrain cholinergic circuits, which are involved in learning, memory and attention. Specifically, following adolescent intermittent ethanol exposure in rodents (AIE), there is a significant suppression of the cholinergic phenotype in neurons within the medial septum/diagonal band (MS/DB), which projects to the hippocampus, and the Nucleus basalis of Meynert complex (NbM) that projects to the cortex.^20,24,25,26^ This AIE-induced dysfunction is a result of an epigenetic suppression of the genes that drive the cholinergic phenotype.^27^ The damages to forebrain cholinergic populations are (is?) age-specific; there is no loss seen when the binge ethanol exposure occurs in adult rodents.^26^ However, despite the AIE-induced loss of cholinergic projection neurons to the hippocampus, <u>behaviorally-evoked</u> acetylcholine (ACh) efflux can be spared in the hippocampus following AIE.^17^ This contrasted with what was/has been observed in the medial PFC: following AIE there is a significant suppression of mPFC ACh efflux activated by spatial exploration.^17^ Subsequently, the frontal cortical cholinergic system may be particularly susceptible to AIE-induced toxicity.

All deep cortical layers receive cholinergic projections that originate from the NbM complex, which includes the NbM, horizontal diagonal band (HDB) and substantia innominata (SI).^28,29^ ACh release in the cortex drives neuronal excitability through altering the presynaptic release of other neurotransmitters (glutamate, GABA) and coordinating the dynamics of local circuits.^30^ There are two modes of ACh transmission, tonic and phasic (transient), and the roles of these two patterns appear to be dissociable. The faster kinetics of cholinergic signaling in the PFC mediates cue detection, including cue-triggered changes in goal-directed behavior.^31,32^ In contrast, the slower and more spatially broader ACh fluctuations are important for arousal, encoding, increasing the signal-to noise ratio.^33^ It has been suggested that the slower cholinergic neuromodulation in the PFC favors staying-on-task staying on task, over switching to an alternative action.^34^

In the current study, we assessed changes in key cholinergic markers in the NbM and OFC, and measured tonic ACh efflux in the orbital frontal cortex (OFC) during a spatial navigation task to determine whether AIE broadly changes cholinergic modulation within the PFC. We assessed behavior in a reversal task to determine if AIE disrupts the ability to change discrete action plans. Finally, we examined whether AIE leads to long-term changes in dendritic architecture within the OFC. We found that AIE resulted in a loss of choline acetyltransferase (ChAT) expression in the NbM and vesicular acetylcholine transporter (VAChT) in the OFC, as well as massively suppressed <u>behaviorally-evoked</u> ACh in the OFC. However, AIE appears to increase apical dendritic complexity in the OFC without influencing spine density or maturity. This increase in dendritic complexity is likely a compensatory reaction in neurotransmitter function following AIE.

## General Experimental Procedures

### Subjects

Pre-adolescent male and female Sprague-Dawley rats were obtained from litters bred at Binghamton University animal facility. Only one pup of each sex (both sexes equally distributed in all treatment conditions) from a litter was randomly assigned to a given treatment condition (see below). All rats were maintained in a temperature (20 degrees) and humidity-controlled colony room in a light/dark cycle (7:00-19:00) and were housed in pairs or, if needed, a triad by sex. Rats in both Treatment conditions (see below) had free access to water and standard rat chow (Purina Lab Diet 5012), except in Experiment 2, where the rats were initially food restricted two weeks prior to the start of behavioral tasks in order to maintain weights at 85% of their free-feeding body weight values.

All procedures were in accordance with the National Institutes of Health (NIH) Guide for Care and Use of Laboratory Animals and the Institutional Animal Care and Use Committee (IACUC) at the State University of New York at Binghamton. The protocols used in the present experiment were approved by the Binghamton University Institutional Animal Care and Use Committee.

### Adolescent Intermittent Ethanol Exposure

Pre-adolescent rats (PD 25) were subject to 16 intragastric gavages of either 20% ethanol (*v/v; AIE; Experiment 1= 7 males, 7 females; Experiment 2= 11 males, 9 females; Experiment 3 =6 males and 6 females*) or water control (CON*; Experiment 1= 6 males, 6 females; Experiment 2= 9 males, 9 females; Experiment 3 = 6 males and 5 females)*, administered at a dose of 5 g/kg. The dosing schedule followed a 2-day on/off cycle, where rats were dosed once per day for 2 days, followed by a 2-day recovery period. Blood samples were collected an hour following the first and eighth gavage, and blood ethanol content (BEC) levels were determined using an AM1 Alcohol Analyzer (Analox Instruments, Ltd, UK). Figure 1 illustrates the timeline and BECs for all experiments.

**Figure 1.**
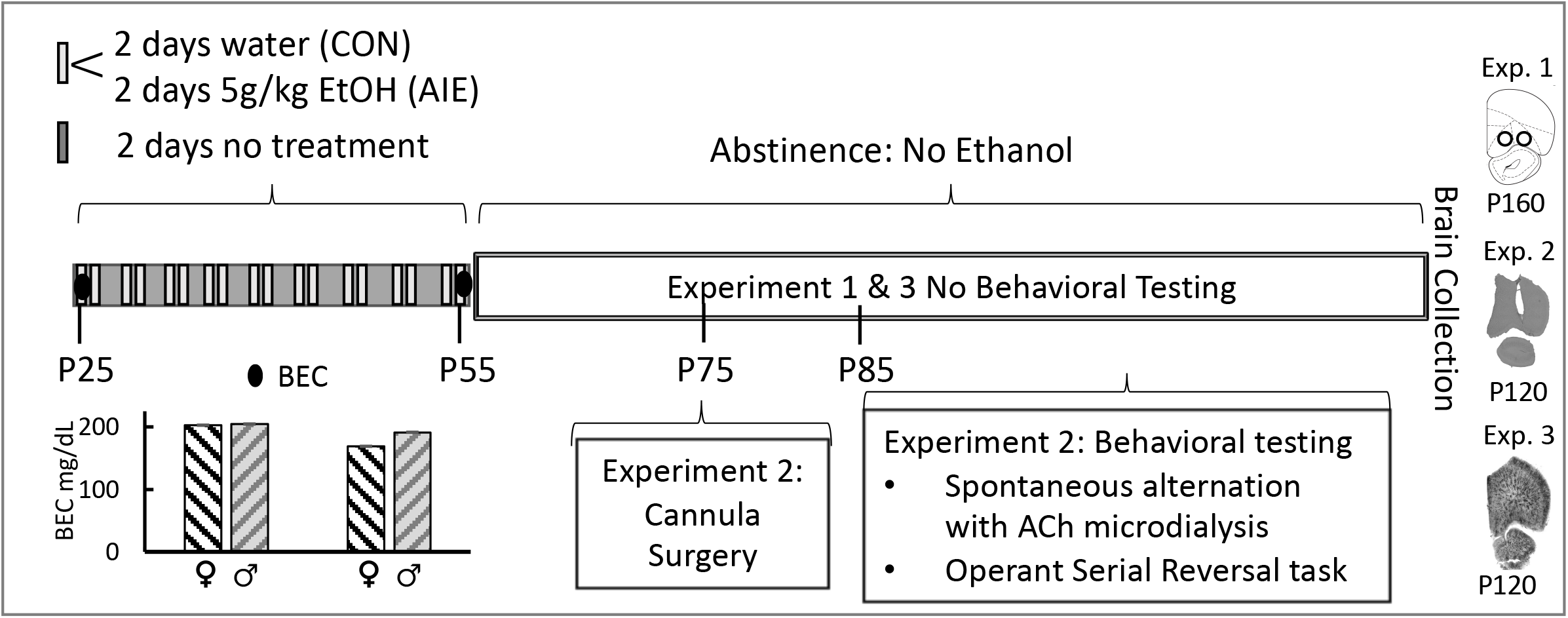
Adolescent Intermittent Ethanol (AIE) exposure protocol and experiment timeline. Schematic outlining the AIE exposure protocol, and the timeline for cannula implantation surgery (Exp 2), behavioral testing (Exp 2) and tissue collection (Exp 1, 2, 3). Blood ethanol concentrations (BECs) following the first and last intraoral gastric gavage are illustrated in the chart insert. BECs did not significantly differ between the sexes.

## Experiment 1

Choline acetyltransferase (ChAT) is responsible for the biosynthesis of acetylcholine and is an indicator of the functional state of cholinergic neurons. Vesicular acetylcholine transporter (VAChT) tightly regulates the release of ACh and is found predominantly in synaptic vesicles at nerve terminals. Western blot procedures were used to assess the persistent loss of ChAT in the NbM and VAChT in the OFC following AIE. Rats were decapitated on approximately PD 180 (due to Covid-19 restrictions), and the brains were flash frozen by brief submersion in cold Methyl Butane, that was maintained on dry ice, and then stored at −80 °C until further use. Brain regions of interest, the NBM and OFC were bilaterally micropunched (1.0–2.0 mm; EMS-Core Sampling Tools, Electron Microscopy Sciences, Hatfield, PA, USA) during the cryostat collection procedure (−20 °C). Brain punches were stored at −80 °C until tissue lysis.

Tissues were homogenized in lysis buffer (1% SDS, 1 mM EDTA, 10 mM Tris) containing protease inhibitors (Halt™ Protease Inhibitor Cocktail, Thermo Scientific, Waltham, MA, USA) and centrifuged at 4°C, 12000g for 30 minutes. Protein concentrations were determined using a bicinchoninic acid method (Pierce, Rockford, IL, USA) and compared to bovine serum albumin standards. 30 μg total protein samples of the OFC and NbM samples were denatured and separated by electrophoresis on Novex™ 8–16% Tris-Glycine sodium dodecyl sulfate polyacrylamide gels (Invitrogen, Carlsbad, CA, USA), transferred to a polyvinylidene difluoride membranes (Invitrogen, Carlsbad, CA, USA), and blocked for one hour in 5% BSA, 0.01% Tween-20 in TBS. Following this, membranes were incubated overnight with a goat antibody against ChAT (NbM=1:500 dilution; Millipore, Burlington, MA, USA; AB144P) and mouse antibody raised against VAChT (OFC= 1:1000 dilution; Millipore, Burlington, MA, USA; ABN100). Blots were then exposed to a peroxidase-conjugated secondary antibody (NbM = mouse anti-goat HRP 1:2000 dilution, Santa Cruz, Dallas, TX, USA; OFC = goat anti-mouse HRP 1:10000 dilution, Thermo Scientific, Waltham, MA, USA) for 1 h and protein levels were detected with enhanced chemiluminescence (Pierce™ ECL Western Blotting Substrate, Thermo Scientific, Waltham, MA, USA). β-actin (Chicken anti β-actin −1:2000 dilution (primary antibody), Origene, Rockville, MD, USA, TA349013; Goat anti-chicken HRP – 1:10000 dilution (Secondary antibody), Abcam, Cambridge, UK; Ab97135) was used for normalization.

### Statistical analysis

For Experiment 1, a two-factor ANOVA (Treatment: AIE vs. CON; Sex: Females vs. Males) was used to assess suppression of the cholinergic markers (ChAT, VAChT) in the NbM and OFC. When significance in the omnibus F test was found, Fisher’s LSD post hoc tests were used to assess the differences between Treatment (AIE vs. CON) and between Sex (Females vs. Males). The SPSS statistical package was used for all analyses and values of p < 0.05 was considered significant.

## Experiment 2

### Cannula Implantation Surgery

For Experiment 3, after AIE, rats underwent cannula implantation surgery and recovery for a 3-week period. A single cannula was placed in the OFC, counterbalanced across hemisphere, 2-3 weeks following the cessation of AIE treatment. Prior to surgery, administration of a ketamine (87.5mg/mL)/xylazine (12.5mg/mL) mixture at a dosage of 1.0 mL/kg was administered (intraperitoneal) as anaesthesia. Rodents were placed into a stereotaxic apparatus (David Kopf Instruments, Tujunga, CA, USA). A guide cannula (5 mm; Synaptech Technology Inc., Marquette, MI, USA) was placed into the OFC coordinates relative to Bregma (Fig 2C) = AP: +4.7, ML: +/−2.0, DV: −4.0 (Paxinos & Watson, 2013). Dental acrylic cement with anchor bone screws secured guide cannula to the skull. Carprofen (5 g/kg; Zoetis, Kalamazoo, MI, USA) was administered prior to surgery, as well as 24-hours and 48-hours post surgery, as an analgesic. After surgery, rats recovered for 10 days before behavioral testing. Spontaneous alternation testing (with ACh microdialysis) occurred first, followed by operant serial reversal learning.

**Figure 2.**
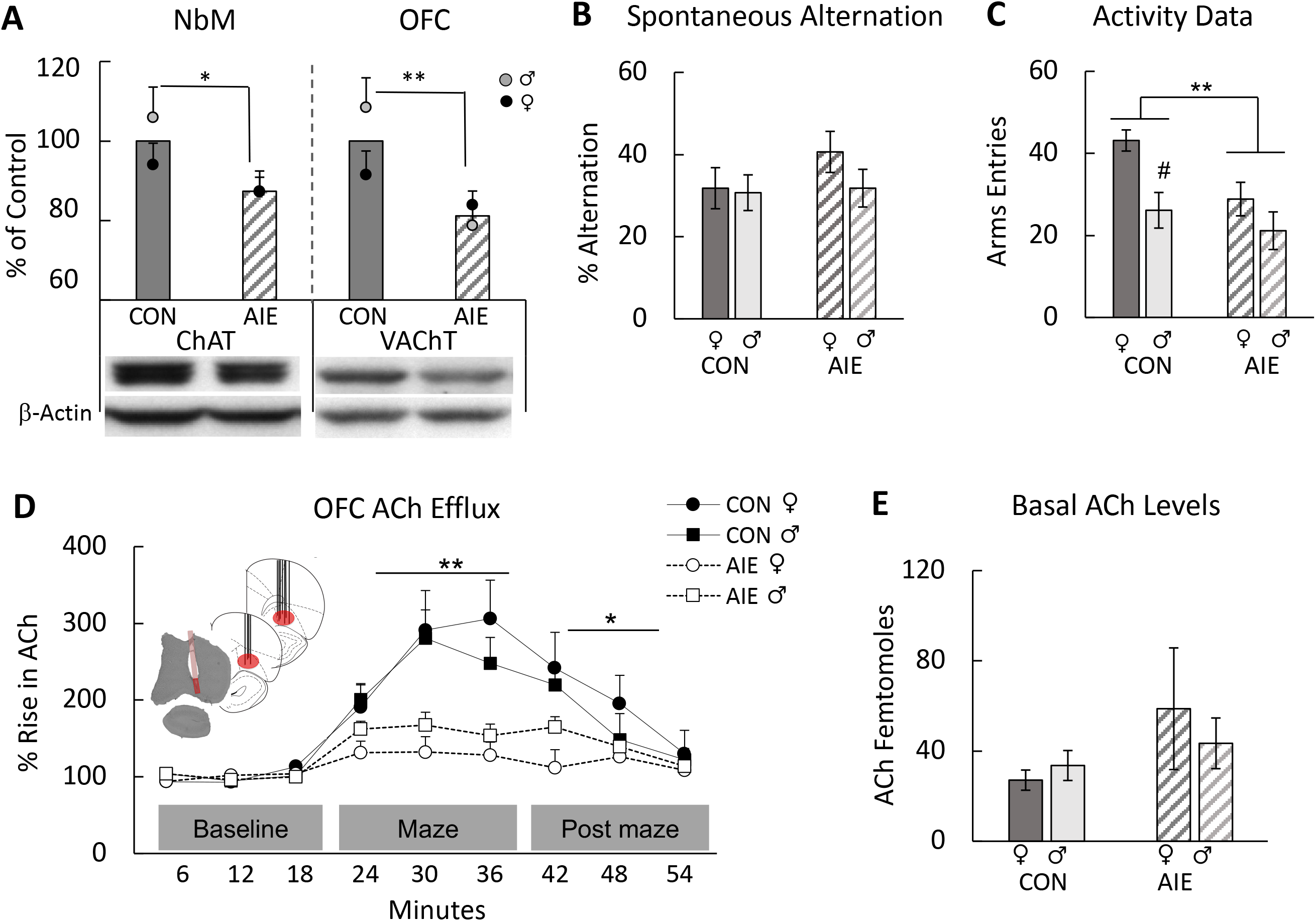
Adolescent Intermittent Ethanol (AIE) leads to a suppression of cholinergic markers in the Nucleus basalis Magnocellularis (NbM)-Orbital Frontal Cortical (OFC) circuit and blunts acetylcholine (ACh) efflux within the OFC during spontaneous alternation, but does not impair spatial exploration. (A) Western blot analyses revealed that AIE decreased the choline acetyltransferase (ChAT) protein levels in the NbM and vesicular acetylcholine transporter (VAChT) protein levels in the OFC, compared to the control condition. Data are presented as the mean ±SEM percent of control. (B) Average percent alternation scores by AIE treatment. There was no effect of AIE on alternation scores. Data are presented as the mean ±SEM. (C) Arm entries during spontaneous alternation. AIE rats were less active than and control rats. In the Control condition, females were more active than males. Data are presented as the mean ±SEM percent. (D) Orbital frontal ACh efflux before (baseline), during (maze) and after spontaneous alternation behavior (after maze). There was a main effect of phase, with ACh levels increasing during maze behavior. There was also a significant interaction between AIE treatment and phase: Regardless of sex, AIE blunted the rise in OFC ACh efflux during maze exploration and into the post maze period. Data are presented as the mean ±SEM percent of baseline. (E) There were no Treatment nor Sex differences in the basal levels of ACh. Data are presented as the mean ±SEM. Treatment effects*= p<0.05, **= p<0.01, # represents a significant Sex effect, *p*< 0.05

### Spontaneous alternation testing with in vivo microdialysis

Each rat was food restricted overnight and tested on a single spontaneous alternation session. Both, ethanol and water pretreated animals, were at similar weights prior to food restriction, and all animals lost weight at similar rates. In vivo microdialysis protocols were followed as previously described.^35^ On the day of testing, a microdialysis probe (S-5020, 2mm; Synaptech Technology Inc., Marquette, MI, USA) was inserted into OFC guide cannula and the rat was placed into an opaque habituation chamber to acclimate for a period of 60-min prior to maze testing. The probe was connected to a CMA microinfusion pump (CMA/400 pump) and an artificial cerebrospinal fluid solution (7.4 pH solution: 127.6 mM NaCl, 0.9 mM NaH_2_PO_4_, 2 mM Na_2_HPO_4_, 4 mM KCl, 1.3 mM CaCl_2_ dihydrate, 1.0 mM glucose, and 0.9 mM MgCl_2_) with 500 nM neostigmine hydrobromide (Sigma-Aldrich Corp., St. Louis, MO, USA) was perfused continuously at a rate of 2.0 μL/min. Baseline dialysate collection began after the 60-minute acclimation period, and samples were collected for three 6-minute intervals.

Spontaneous alternation was conducted in a plus maze (105.5 cm × 14.4 cm × 15 cm). The rat was placed into the center of the apparatus and allowed to explore the maze for 18-minutes of testing, during which arm entries (all four paws within an arm) were recorded. Dialysate was continuously collected during maze behavior. An alternation was defined as entry into four different arms in overlapping successive sequences of 4 arm entries (for example, in the successive arm entries of B, A, D, C, A, D, C, A, D, B, C, D, B, A; the first sequence of BADC was an alternation, but the next 4- arm sequence ADCA was not). The percent alternation score is equal to the ratio of actual alternations to possible alternations (total alternations/[trial number-3] X 100). Following maze testing, the rat was returned to the opaque habituation chamber for an 18-minute collection of post maze dialysate.

### High-performance liquid chromatography

High-performance liquid chromatography (HPLC) with electrochemical detection (Amuza, San Diego, CA, USA) was used to assay ACh from dialysate samples. Chromatographs obtained were analyzed using the software program Envision (provided by Amuza, San Diego, CA, USA). ACh peaks were quantified by comparison to peak heights of standard solutions (100 nM, 20 nM, and 4 nM standards).

### Operant Serial Reversal Methods

Operant chambers (30 cm × 33 cm × 23 cm; Med Associates Inc., St. Albans, VT, USA) were enclosed within sound attenuating boxes (59 cm × 55 cm × 36 cm) equipped with a running fan. Each operant chamber contained two retractable levers separated by a magazine which dispensed food reward (Rodent Purified Dustless Precision Pellet; Bio-Serve, Flemington, NJ, USA). Above each retractable lever were stimulus lights, and a house light located on the opposite side of the chamber from the magazine provided the main source of illumination during operant training. Operant chambers were interfaced with MED-PC (Med Associates Inc. St Albans, VT, USA)

Rats were gradually food restricted to 85% of their free feed weight over the course of one week before the start of pre-training, and this level of food restriction was maintained throughout operant testing. Each of the 2 days prior to the start of pre-training, rats were exposed to the food reward used during operant training. During operant pre-training and training, rats were assigned to operant chambers and underwent testing on one task per day, for a maximum of 1-hour duration of training.

During pre-training, rats were placed in the operant chamber and randomly assigned to be shaped in pressing the left or right lever until 50 lever presses were recorded within a 30-minute period. Only one lever was extended during this task while both cue lights were illuminated. Following mastery of the left or right lever shaping, rats were trained to press the opposite lever under the previously mentioned conditions until criterion was met. Each response was rewarded on a Fixed-Ratio-1 reward schedule. After shaping left and right lever presses, rats underwent 90 trials of retractable lever training. During each trial of the retractable lever training either the left or right retractable levers randomly extended for 10 seconds, while both cue lights were illuminated. Rats were rewarded following a lever press within this 10 second window, otherwise the trial was considered an omission. In order to master the retractable lever training task, rats were required to make fewer than 5 omissions in 90 trials over two consecutive days.

The spatial reversal task had an initial acquisition phase which was followed by three rule shifts. For the acquisition task, the rewarded (left or right) lever was opposite of the rats predetermined side bias. A randomly determined cue light above a lever was illuminated for 3 seconds prior to lever extension, remained illuminated until the end of the trial, and was not predictive of the rewarded lever. Both levers were presented and the rat was required to make a choice within 10 seconds. A correct response dispensed a single food pellet, while incorrect responses and omissions (not making a choice within 10 seconds) were not rewarded. After each response, or omission, a 20 second inter-trial-interval preceded the start of the next trial. For all tasks, 10 consecutive correct responses were required to complete the task. If the rat did not complete the task in 200 trials over the course of 1 hour, the animal would be retested on the same task the next day and responses on the task would be combined. The next three reversal tasks required the rat press the opposite lever from the previous phase. The number of trials required to reach criterion (10 consecutive correct lever presses), the number of errors (unrewarded lever presses), the latency to lever press, and the latency to collect food reward were recorded for each phase.

### Tissue Collection

One week after completion of the operant set shifting test, rats were euthanized (Fatal-Plus, Vortech Pharmaceuticals, Dearborn, MO, USA) by rapid decapitation. The whole brain was removed, cut sagittal and placed in a 4% paraformaldehyde solution for 1 week followed by submersion into a 30% sucrose solution in 0.1 M PBS at 4 C for 4-8 days.

### Cresyl violet staining

To determine the location of cannula implantation and probe insertion, the hemispheres brains were sliced at 40 microns using a microtome (Sm2000r Leica Biosystems, Wetzler, Germany) and stored in cyroprotectant solution (62.8 mg Na2HPO4, 160 mL dH2O, 120 mL 120 mL glycerol) at −20 C.

Brain sections from the OFC were mounted onto slides immediately after slicing and stained using a cresyl violet protocol.^25,36^ Slides were placed in a series of ethanol (95%, 70%, 50%) and water (distilled H20) treatments to rehydrate the material before staining with the cresyl violet stain solution (FD Cresyl Violet Solution; PS102-2; FD Neurotechnologies, Columbia, MD, USA). The tissue was stained in cresyl violet for 5 min and dehydrated again in reverse order with placing the tissue in distilled water, followed by 50%, 70% and 95% ethanol and finally Citrisolv (04-355-121; Fisher Scientific, Waltham, MA, USA). Slides were cover slipped using Permount mounting medium (SP15-500; Fisher Scientific, Waltham, MA, USA) in preparation to be microscopically analyzed.

### Statistical analysis

A repeated-measures analysis of variance (ANOVA), with Sex (AIE female × AIE male) as the between-subjects factor, was used to analyze BEC (across time). For Experiment 2, a two-factor ANOVA (Treatment: AIE vs. CON; Sex: Females vs. Males) was used to assess the spontaneous alternation data such as percentage of alternation and number of arms entries. To identify changes in ACh efflux from baseline, during maze, and after behavioral testing phases, the microdialysis data were analyzed using a two-factor ANOVA with repeated measures (Phase × Sample). Fisher’s LSD post hoc tests were used to assess the differences between Treatment (AIE vs. CON) and between Sex (Females vs. Males). Operant set shifting data were analyzed using two-factor ANOVA for each dependent variable in each task. The SPSS statistical package was used for all analyses and values of p < 0.05 was considered significant.

## Experiment 3

### Golgi-Cox staining

Approximately 60 days after treatment (PD115-125), rats (AIE, CON) were euthanized (Fatal-Plus, Vortech Pharmaceuticals, Dearborn, MO, USA) by rapid decapitation under light. The whole brain was removed, blocked coronally for the orbital frontal cortex and placed in Golgi-Cox solution A + B (FD Rapid GolgiStain Kit; Cat.#PK401, FD Neuro Technolgies, Inc. Columbia, MD, USA) for 2 weeks, followed by Solution C for one month at room temperature in the dark. The first block was sliced coronally at 200 μm-thick using a sliding microtome (Sm2000r Leica Biosystems, Wetzler, Germany). Slices (Bregma 5.16 to 3.72)^37^ were immediately transferred sequentially to double gelatinized slides at 6 slices per slide using Solution C. After being kept in the dark for maximum 72-hours, the slices were stained under red light with solution D+E and underwent a dehydration process according with the protocol as outlined by FD Rapid GolgiStain Kit User Manual. Slides were cover slipped using Permount immediately after development. Following cover slipping, the slides were left untouched in a dark room for one month until completely dry.

### Dendritic Analysis

Prior to analysis, the slides were coded randomly by a secondary experimenter to blind the experimenter to condition. Using a computer-based neuron tracing system (NeuroLucida; MicroBrightField, Williston, VT, USA), eight neurons for each animal in the OFC were identified under the 5x objective. Using a 20x objective, the experimenter ensured that the neuron fit the following criteria: pyramidal-shaped cell body, located in layer III in the OFC, apical dendrite with depths between 200 and 500 μm from the cortical surface, have branch orders 3 and 5 with at least 60 μm each on the apical dendrite and neurons was not broken. Once the neuron had been determined to meet the aforementioned criteria, the experimenter traced the neuron’s cell body and apical dendrites, under the 40x objective. In order to determine dendritic complexity, the total length and a Sholl analysis were performed for each neuron.

For all neurons, spine analysis was performed on the same dendritic branches for Order 3 and Order 5 previously traced using the 100x magnification with an oil immersion lens. The spines were assessed and marked in a total of 60μm in length of branches in orders 3 and 5 of the apical dendrite and the spine density was calculated per 10μm. All spines identified were also classified as mature and immature following the protocol outlined by Risher and collegues.^38^ The percentage of mature and immature spines were calculated for each animal.

### Statistical analysis

The effect of Treatment and Sex on apical dendritic length were analyzed using a two-factor analysis of variance (ANOVA). A two-factor ANOVA with repeated measures for Distance from the soma was used in Sholl analysis, followed by Fisher’s LSD post hoc tests to assess the differences in dendritic complexity between Treatment (CON vs. AIE) and Sex (Males vs. Females). A repeated measures ANOVA was also used to evaluate within-subject effects (Phenotype [mature, immature] and branch Order [3^rd^, 5^th^]) and between-subject effects (Treatment, Sex) for spine density. Two-factor repeated-measures analysis of variance (ANOVA) with branch Order as the within-subject variable and Treatment and Sex as the between-subject variables was used to analyze the spine density of each spine phenotype separate and the percentage of mature or immature spines. The percentage of mature spines versus immature spines was analyzed using a repeated-measures ANOVA with Phenotype as the within-subject variable. The SPSS statistical package was used for all analyses and values of p < 0.05 was considered significant.

## Results

### AIE rats reached heavy binge-like levels of intoxication

Rats exposed to AIE had high BEC that well exceeded binge EtOH benchmarks of 80 mg/dL^39^ and would be considered extreme binge levels.^40^ There was a significant Experiment X Sample Time interaction (F [2, 34] = 3.99, p < 0.05). In Experiment 2 (first day = 203.73 mg/dl, SEM = 10.85; final day =180.14 mg/dl, SEM = 9.44) and Experiment 3 (first day = 216.31 mg/dl, SEM =12.29; final day = 180.74 mg/dl, SEM = 10.69) the BEC was higher on the first day of gavage compared to the last day of gavage. However, in Experiment 1 there was no difference in BEC as a function of sampling time (First day = 230.46 mg/dl, SEM = 10.51; final day = 250.63 mg/dl, SEM = 9.14). Male and females did not differ significantly in terms of BEC (F [1,34] = 3.45, p = 0.72), and there were no Sex × Experiment interactions detected (F [2,34] = 2.44, p > 0.103).

## Experiment 1

### Cholinergic markers are persistently down regulated in the Nucleus basalis Magnocellularis and the Orbitofrontal cortex

As shown in Figure 2A, ChAT expression within the in the NbM was significantly lower in AIE treated rats (F [1,22] = 5.39, p < 0.05), compared to control rats, and no Sex, or Treatment × Sex interactions were significant. Additionally, VAChT expression in the OFC was significantly reduced in AIE treated rats (F [1, 22] = 9.57, p = 0.005), compared to control rats. No Sex, or Treatment × Sex interactions were detected.

## Experiment 2

### AIE spared spatial memory assessed using a spontaneous alternation task

Analysis of percentage alternation during this task of spatial working memory did not show effect of Treatment or Sex (both: F [1,29] ≤ 1.04, p = 0.32; Fig 2B). However, the number of arm entries during testing were affected by Treatment (F [1,25] =8.74, p = 0.007) and by Sex (F [1,25] = 14.40, p = 0.001; Fig 2C). Fisher’s LSD post hoc tests showed that activity of CON animals in the maze were higher than AIE rats (p=0.007), and females were more active than male rats (p=0.001) during the testing. T-tests showed that CON female rats made significantly more arm entries than CON male rats (t [13] = 4.03; p = 0.001) and more than AIE female rats (t [10] = 2.95; p = 0.014). AIE male rats did not differ from CON male rats (t[13]= 1.15; p = 0.26) or AIE female rats (t[10]= −1.57; p = 0.14). Thus, this type of spatial memory is spared following AIE.

### AIE reduced ACh efflux in the orbital frontal cortex

The analysis of ACh efflux in the OFC indicated a main effect of Treatment (F [1,23] = 12.26, p = 0.002; Fig. 2D). However, there was no effect of Sex or interaction between Treatment and Sex (F [1,23] = 1.23, p = 0.28). While CON rats showed a large increase in efflux of ACh during maze testing, this effect was notably blunted in AIE rats as there were significant interactions between Treatment × Phase (F [2,46] = 10.62, p <0.0001), and Treatment × phase × samples (F [4,92] = 8.11, p < 0.0001). Due to the significant interactions, the ACh efflux data were analyzed separately during behavioral testing (maze) and after maze phases. During maze and after maze phase, AIE-treated rats had blunted OFC ACh efflux compared to CON rats (maze: F[1,23] = 17.02; p < 0.0001; after maze: F[1,23] = 4.53; p = 0.044). Analysis of basal femtomoles ACh levels in the OFC (Fig. 2E) did not show significant effects of treatment (F [1,23] = 2.23, p = 0.14) or sex (F [1,23] = 0.10, p = 0.75).

### AIE did not disrupt operant reversal learning

Figure 3, displays that AIE did not affect the number of trials to criterion or number of errors in any phase of the operant testing. During the acquisition, there was no effect of Treatment or Sex (both F’s[1,34] < 1.0, p > 0.15). There were no group differences in terms of errors on this task as a function of Treatment or Sex (both F’s [1,33] < 1.93, p > 0.15). Latency to lever press was also analyzed and identified no Treatment or Sex effects (all F’s [1,33] < 2.47, p > 0.10). Additionally, no Treatment or Sex effects were present in latency to collect reward (both F’s [1,33] < 2.82, p > 0.10).

**Figure 3.**
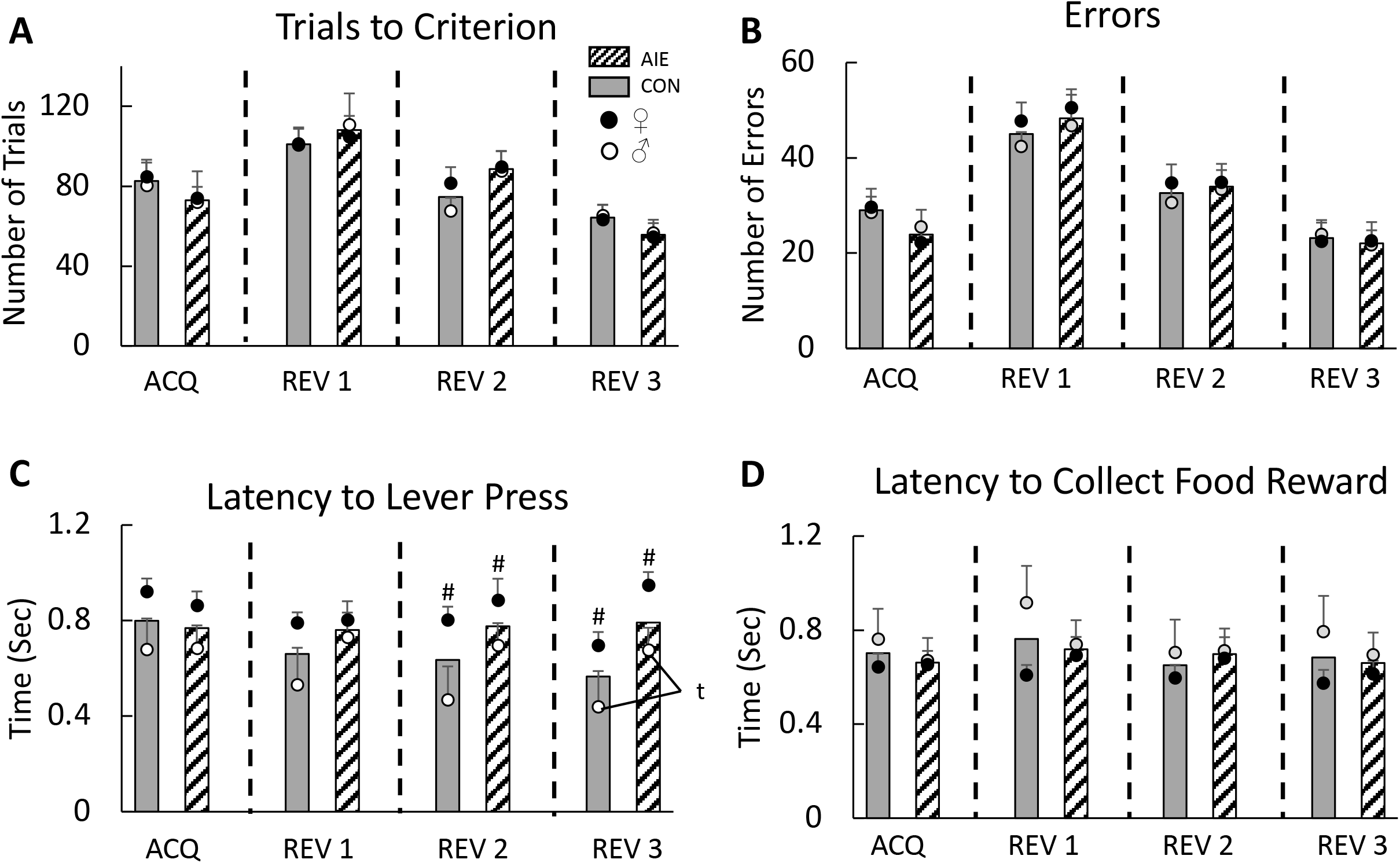
Adolescent intermittent ethanol exposure (AIE) did not affect acquisition, reversal learning nor response times in an operant two-lever paradigm. The number trials to reach criteria (A) and the number of errors made (B) by AIE- and CON-treated female and male rats during acquisition and several serial reversals were not different. The latency to lever press on any task was not affected by AIE (C). However, during the second and third reversal female rats took longer to make a decision than male rats. In addition, there was a trend for male AIE rats to take longer to press the lever then male control rats. Latencies to collect reward (D) were not different as a function of task, sex or AIE treatment. All data are presented as the mean ±SEM. # represents a significant Sex effect, *p*< 0.05; t represents a trend towards significance, p=0.06

For trials to criterion on the first reversal, no Treatment or or Sex differences were observed (both F’s [1,35] <1.0, p > 0.15). Similarly, analyzing the number of errors revealed no Treatment or Sex effects (F’s [1,35) <1.0, p>0.15. Latency to lever press did not differ as a function of Treatment or Sex (both F’s [1,35] <1.59, p > 0.15). Additionally, there were no group differences in latency to collect reward based on Treatment or Sex (both F’s [1,35] < 2.82, p > 0.10).

On the second reversal, no Treatment or Sex differences were observed in the number of trials required to reach criterion (both F’s [1,35] < 2.71, p > 0.10). Additionally, there were no Treatment or Sex differences on the number of errors made (F’s [1,35] < 1.0, p > 0.15). While no Treatment (F [1,35] = 1.62, p > 0.15), differences were observed in latency to lever press, a significant effect of Sex was observed (F [1,35] = 4.67, p < 0.05): Male rats (M = 0.59 seconds; SEM = 0.08) responded more quickly to lever presentation than female rats (M =0.84 seconds; SEM = 0.09). Lastly, no Treatment or Sex differences were identified in latency to collect food reward (F’s [1,35] < 1.0, p > 0.10).

On the third, and final, reversal no Treatment or Sex effects were observed in trials or errors to reach criterion (both F’s [1,35] < 1.67, p > 0.10). Analyzing latency to lever press on the third reversal revealed a trending Treatment effect (F [1,35] = 3.74, p = 0.06), where CON rats (M = 0.56 seconds, SEM = 0.09) responded quicker to lever presentation than AIE rats (M = 0.81 seconds, SEM = 0.09). There was also a significant main effect of Sex F (1,35) = 4.34, p < 0.05, with female rats (M = 0.82 seconds, SEM = 0.09) responding slower than male rats (M = 0.57 seconds, SEM = 0.09). Lastly, when examining latency to collect food reward, there were no differences as a function of Treatment or Sex (both F’s [1,35] < 2.31, p > 0.10).

## Experiment 3

Representative images and tracing of a neuron cell body, apical dendrite and dendritic spines in OFC are provided in Fig 4A.

**Figure 4.**
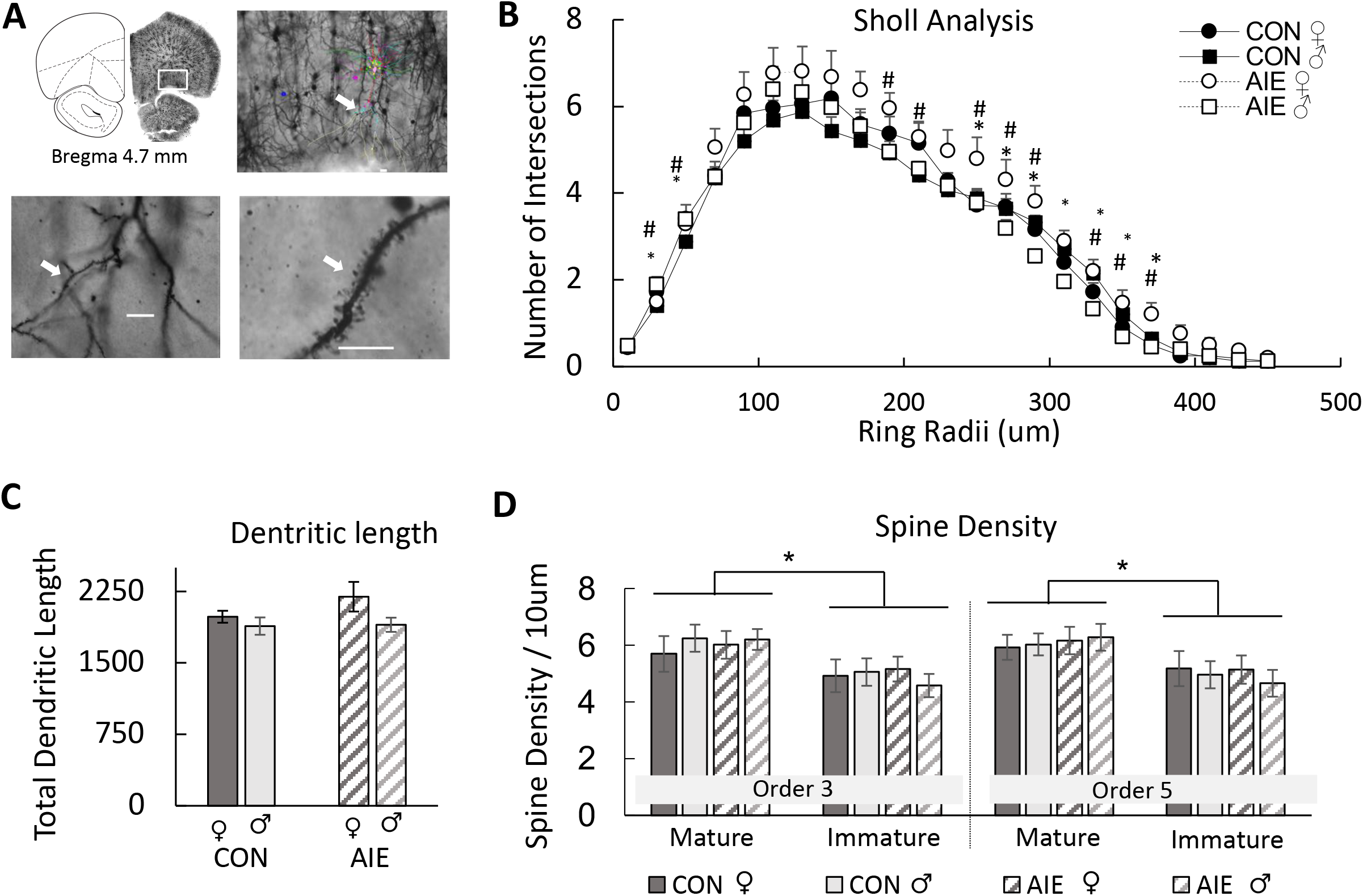
Adolescent intermittent ethanol exposure increased the apical dendritic complexity in pyramidal neurons within the orbital frontal cortex (OFC). (A) Representative image of the apical dendrites of pyramidal neurons in the OFC (top right; 20x objective; Scale bar = 25 μm) in OFC. The tracing demonstrates the ordered branches and the arrow shows the order 5. Bottom images show high magnification images of spines on order 5 apical dendritic branches. Bottom left: magnified (40x objective) image of apical branches from a neuron in the OFC. The right image is then magnified to a 100x objective (oil lens) in order to determine the spine density and phenotype (Scale bar = 10 μm). (B) Scholl analysis revealed that AIE increased the number of dendritic branches, close to the soma for males and away from the soma for females. (C) Dendritic length was not affected by AIE. (D) Spine density was affected by AIE. However, there were more spines of the mature phenotype, relative to the immature phenotype, in both 3 and 5 order branches. All data are presented as the mean ±SEM. # represents a significant Sex effect, *p*< 0.05; * represents significant Treatment (CON vs. AIE) effect, *p*< 0.05; #* together represents a Sex X Treatment interaction *p*< 0.05

### AIE and Sex influence apical dendritic complexity in the OFC

Sholl analysis was used to analyze dendritic complexity. The average number of intersections with each radii 20μm of concentric sphere surface was analyzed for each group (Fig 4B). A repeated measure ANOVA (Treatment × Sex × Distance from soma) of apical dendritic intersections in OFC revealed main effects of Treatment (F [1, 16] = 8.35, p = 0.011), Sex (F [1, 16] = 17.67, p = 0.001) and Distance from soma (F [23, 368] = 35.86, p < 0.001). Also, the analysis revealed several significant interactions: Treatment X Sex (F [1, 16] = 5.48, p = 0.032), Treatment X Distance (F [23, 368] = 1.73, p < 0.021) and Distance X Sex (F [23, 368] = 1.86, p < 0.01). Fisher’s LSD post hoc tests showed that female rats have significantly more intersections than male rats, particularly at the later intersections (190 μm [p = 0.028], 210 μm [p = 0.041], 270 μm [p = 0.049], 290 μm [p = 0.016]).

A detailed analysis of the influence of AIE on dendritic complexity as a function of sex revealed that the dendritic complexity of CON male rats started with less intersections than AIE male rats at the early intersections (30 μm (t [10] = 2.32, p = 0.043) and 50 μm (t [10] = 2.57, p = 0.028); however, at the later intersections (290 μm through 350 μm (t’s [10] > 4.62, p’s < 0.01), the CON male rats have significantly more intersections than the AIE male rats. There was a trend for CON female rats to have fewer intersections compared with AIE female rats at the later intersections (370 μm (t [9] = 2.16, p = 0.058) and at 390 μm (t (9) = 2.19, p = 0.055). AIE female rats had more intersections than AIE male rats at 270 μm through 390 μm (t [10] < 3.50, p < 0.05).

### AIE did not affect the length of OFC dendrites

Total dendritic length is another dendritic complexity parameter that can be used to examine the effects of AIE as a function of sex (Fig 4C). A two-way ANOVA showed only a trend for female rats to have a longer dendritic length in apical dendrites than male rats (F [1, 22] = 3.43, p = 0.079), regardless of treatment condition. There was no effect of Treatment on the total dendritic length of apical dendrites in the OFC.

### AIE exposure did not alter the total spine density or alter the density of mature or immature phenotypes, but mature dendritic spines were the most prevalent in the OFC

The ANOVA revealed no effects of Order of branches (3 vs 5; F[1,19] = 0.40, p = 0.53), Treatment (F[1,19] = 0.012, p = 0.91) or Sex (F[1,19] = 0.013, p = 0.91) in total spine density. However, the analysis revealed a main effect of Phenotype (F [1, 19] = 7.067, p = 0.016, see Fig 4D): There was a significantly higher percentage of mature spines, relative to immature spines, regardless of Treatment or Sex.

In a separate analysis, we analyzed the density of immature and mature dendritic spines. Mushroom and stubby phenotypes were classified as mature spines, whereas filopodia and thin phenotypes were classified as immature spines. There was a trend for a significant interaction between Order, Treatment and Sex on the density of stubby spines (F [1, 19] = 3.849, p = 0.065). Overall, this interaction is driven by a small decrease in the density of stubby spines in Order 5 branches (average of spine density in Order 5 = 1.21 and Order 3 = 1.29), in females (average of spine density in females = 1.09 and males = 1.41) and AIE rats (average of spine density in AIE = 1.20 and CON=1.30). Furthermore, there were no main effects of Order (F [1,19] = 0.40, p = 0.83), Treatment (F [1,19] = 0.15, p = 0.69) or Sex (F [1,19] = 0.43, p = 0.51) on the percentage of mature or immature spines.

## Discussion

The key findings of these experiments are the following: (1) Regardless of sex, there is persistent loss of ChAT protein expression in the NbM, and a loss of VAChT protein expression in the OFC, demonstrating long-term disruption of this cholinergic circuit following adolescent ethanol exposure; (2) Independent of sex, behaviorally-evoked cholinergic tone in the OFC is dramatically and persistently disrupted by binge-type ethanol exposure during adolescence; (3) The branching of apical dendric trees in the OFC is persistently increased following binge-type ethanol exposure during adolescence, but sex influences whether the increase in branching occurs near or away from the soma; (4) Not all types of reversal learning are impaired by adolescent binge-type ethanol exposure.

Previous work demonstrated that there is about a 30% suppression of the cholinergic phenotype in the nucleus basalis magnocellularis complex.^25,26^ This nucleus innervates the cortical mantle, including the OFC. ^28,29^ The results from Experiment 1 demonstrate the loss of the expression of ChAT protein in the NbM and revealed a related loss of VAChT in the OFC. A decrease in VAChT expression influences the amount of ACh loaded in synaptic vesicles and therefore its release.^41^ In adult mice, genetic elimination of VAChT from the forebrain caused deficits in reversal learning assessed by the Morris water maze and operant two-choice touch screen.^42,43^ Furthermore, performance on a 5-choice serial reaction time task is impaired by forebrain VAChT deletion, while impulsive and perseveration behaviors are spared.^42^

It should be noted that even small to moderate changes in the expression of VAChT has the potential to change synaptic transmission.^41^ Thus, although we do not see a general suppression of ACh levels, as basal femtomole values of ACh are not changed by AIE, the behaviorally-evoked changes in cholinergic tone are dramatically suppressed in the OFC following AIE. Interesting, hippocampal ACh efflux during maze mapping is not affected by AIE, but medial PFC ACh levels are also suppressed.^25^ Thus, despite the reduction in neurons expressing the cholinergic phenotype in all basal forebrain nuclei following AIE^26^, only the cortical projection regions display a significant reduction of activity-dependent cholinergic tone. Thus, the forebrain cholinergic projection to PFC is particularly vulnerable to binge-type adolescent ethanol exposure.

We did not see impairment of spatial mapping on the spontaneous alternation task and this is supported by past work.^17^ However, we did see that AIE rats were less active on the maze, potentially due to their known increase in anxiety.^44,45^ There were no sex driven effects, nor interactions with AIE, on cholinergic parameters or spatial maze performance.

An assessment of reversal learning was included as performance on such tasks can be dependent on the OFC^46,47^ and reversal learning can be disrupted by AIE.^12,27,48^ However, the serial reversal paradigm used did not reveal AIE-induced deficits. The operant two-choice lever paradigm, without modulating reinforcement values, may have not been complex enough to evoke the OFC as subjects may have adapted a simple win-shift strategy rather than a complex stimulus, response, reward contingency map, which is OFC dependent.^46,47^

The persistent microstructural changes to the OFC induced by intermittent ethanol exposure throughout adolescence are somewhat subtle: an increase in dendric branch complexity at different distances from the soma that is modulated by sex. Male rats exposed to AIE had an increase in dendritic branching close to the soma, whereas female rats exposed to AIE had an increase in dendric branching away from the soma. Abnormal dendritic branching can lead to decreased or excessive synaptic connectivity and can contribute to alterations in cognitive functions.

Dendritic arborization is dynamic and determines the synaptic input field, but itself is influenced by synaptic activity and pathological conditions can lead to dendritic remodeling.^49^ As a morphogen, ACh has powerful effects on dendritic arborization in adulthood and development.^50–52^ Release of ACh can reconfigure cortical microcircuitry. However, various dendritic changes are region specific in response to cholinergic modulation.^50–51^ The AIE-induced altered ACh tone is unlikely to be the sole contributor to dendritic remodeling following chronic ethanol exposure during adolescence or adulthood. Other studies have examined PFC dendritic remodeling after ethanol exposure on a shorter time scale and found that withdrawal is a critical component. A recent review demonstrates that cycles of alcohol intoxication and withdrawal lead to poorer behavioral and brain outcomes, especially if the ethanol exposure occurs during development.^53^

In rodents, ethanol exposure, followed by withdrawal, changes dendritic arborization and dendritic spine density in the PFC and hippocampus.^38,54-56^ Excitotoxic events, such as withdrawal from ethanol, can cause dendritic remodeling; For example 3-hours following Chronic Intermittent Ethanol (CIE) in adult mice, but not during intoxication, there was an increase the number of dendritic intersections at shorter distances from the soma in both basal and apical dendrites within the mPFC.^55^ This study also found CIE increased the spine density of pyramidal neurons in the mPFC, but CIE did not alter the total combined length of the apical and basal dendritic trees.

Similarly, in the lateral OFC, CIE in the absence of withdrawal in adult mice, had no effect on spine morphology or spine density in layer II/III pyramidal cells. However, when mice were allowed to undergo a 7-day withdrawal period, there were significant increases in spine density and long thin spines.^57^ The mature spine phenotype, mushroom-shaped or short stubby spines, were unchanged by CIE.

Different PFC regions show differential reactions to ethanol exposure and withdrawal. The effects of adult chronic ethanol exposure on the length of basal and apical dendrites of lateral OFC and medial PFC neurons were examined 3-days following ethanol withdrawal.^58^ It was found that following 3-days withdrawal from CIE there was hypertrophy of distal apical non-terminal dendrites and dendritic retraction of proximal apical dendrites of prelimbic (PrL) pyramidal neurons, but no effect on the arborization of pyramidal neurons in the OFC or infralimbic (Il) cortex. Adult CIE increases dendritic arborization and spine densities within basal and apical dendrites of pyramidal neurons represents an aberrant reorganization.^55^

Intermittent ethanol exposure during the adolescent period, relative to the adult period, suppresses spine density in a region-specific manner within the mPFC. The spine density within IL was significantly reduced 3 days following intermittent ethanol exposure, relative to air, in the adolescent exposed group, but not in adult-exposed mice.^54^ In contrast, in the PrL, overall spine density was not affected by AIE exposure, but was significantly lower in the adult mice than the adolescent mice. However, rats exposed to intermittent ethanol during early-mid adolescence (PD28–42), with brains assessed in adulthood, displayed a persistent increase in the density of long/thin dendritic spines, with no change in stubby or mushroom spines of layer 5 pyramidal neurons in the PrL cortex.^56^ Thus, the cortical microstructure recovery after protracted abstinence may also be region specific.

In summary, repeated extreme binge-type alcohol exposure leads to a long-term persistent change in the OFC cholinergic profile and microstructural organization. The dramatic suppression of behavioral-evoked ACh may contribute to the increase in dendritic complexity, as ACh is a known morphogen. The slight increase in apical dendritic branching within the OFC may be a compensatory response to the reduced cholinergic tone and altered cortical dopamine levels^56^, both reported following AIE. Data from human imaging and neuropsychological profile assessment suggests that there are persistent effects of adolescent binge drinking on specific cognitive domains and <u>frontcortical</u> measures that extend well into adulthood, and this has been replicated in rodents.^59^ However, the nature and degree of persistent brain changes and cognitive dysfunction is still being determined. Behavioral tests that have a more complex integration of changes in environmental and reward continencies contingencies? with the need to modify response strategies are likely to be more sensitive to AIE-induced brain damage within the OFC.

## Abbreviation List

ACh: Acetylcholine
AIE: Adolescent intermittent ethanol
ChAT: Choline acetyltransferase
CON: Control gavage treatment
HDB: horizontal diagonal band
MS/DB: medial septum/diagonal band
NbM: Nucleus basalis of Meynert
OFC: Orbital frontal cortex
PFC: prefrontal cortex
SI: substantia innominata
VAChT: Vesicular acetylcholine transporter

## Acknowledgements

None

## Notes

Conflict of interests: No

Funding Sources: This work was supported by U01 AA028710 (LMS), P50 AA017823 (LMS), T32 AA025606 (JDJ)

### Competing Interest Statement

The authors have declared no competing interest.

## References

1. Broadwater MA, Lee SH, Yu Y, Zhu H, Crews FT, Robinson DL, Shih YI. Adolescent alcohol exposure decreases frontostriatal resting-state functional connectivity in adulthood. Addict Biol. 2018;23:810–823.

2. Squeglia LM, Tapert SF, Sullivan EV, Jacobus J, Meloy MJ, Rohlfing T, Pfefferbaum A. Brain development in heavy-drinking adolescents. Am J Psychiatry. 2015;172:531–42.

3. Silveri MM, Dager AD, Cohen-Gilbert JE, Sneider JT. Neurobiological signatures associated with alcohol and drug use in the human adolescent brain. Neurosci Biobehav Rev. 2016;70:244–259.

4. Spear LP. Effects of adolescent alcohol consumption on the brain and behaviour. Nat Rev Neurosci. 2018;19:197–214.

5. Giedd JN, Rapoport JL. Structural MRI of pediatric brain development: What have we learned and where are we going? Neuron. 2010;67:728–734.

6. Herting MM, Sowell ER. Puberty and structural brain development in humans. Front Neuroendocrinol. 2017;44:122–137.

7. Sowell ER, Delis D, Stiles J, Jernigan TL. Improved memory functioning and frontal lobe maturation between childhood and adolescence: a structural MRI study. J Int Neuropsychol Soc. 2001;7:312–22.

8. Stiles J, Jernigan TL. The basics of brain development. Neuropsychol Rev. 2010;20:327–348.

9. Jadhav KS, Boutrel B. Prefrontal cortex development and emergence of self-regulatory competence: The two cardinal features of adolescence disrupted in context of alcohol abuse. Eur J Neurosci. 2019;50:2274–2281.

10. Sanhueza C, García-Moreno LM, Expósito J. Weekend alcoholism in youth and neurocognitive aging. Psicothema. 2011;23:209–214.

11. Crews FT, Nixon K. Mechanisms of neurodegeneration and regeneration in alcoholism. Alcohol. 2009;44:115–127.

12. Vetreno RP, Crews FT. Adolescent binge drinking increases expression of the danger signal receptor agonist HMGB1 and Toll-like receptors in the adult prefrontal cortex. Neuroscience. 2012;226:475–488.

13. Coleman LG, Liu W, Oguz I, Styner M, Crews FT. Adolescent binge ethanol treatment alters adult brain regional volumes, cortical extracellular matrix protein and behavioral flexibility. Pharmacol Biochem Behav. 2014;116:142–151.

14. Vargas WM, Bengston L, Gilpin NW, Whitcomb BW, Richardson HN. Alcohol binge drinking during adolescence or dependence during adulthood reduces prefrontal myelin in male rats. J Neurosci. 2014;34:14777–14782.

15. Liu W, Crews FT. Adolescent intermittent ethanol exposure enhances ethanol activation of the nucleus accumbens while blunting the prefrontal cortex responses in adult rat. Neuroscience. 2015:293;92–108.

16. Gass JT, Glen WB, Jr., McGonigal JT, Trantham-Davidson H, Lopez MF, Randall PK, Yaxley R, Floresco SB, Chandler LJ. Adolescent alcohol exposure reduces behavioral flexibility, promotes disinhibition, and increases resistance to extinction of ethanol self-administration in adulthood. Neuropsychopharmacology. 2014;39: 2570–2583.

17. Fernandez GM, Lew BJ, Vedder LC, Savage LM. Chronic intermittent ethanol exposure leads to alterations in brain-derived neurotrophic factor within the frontal cortex and impaired behavioral flexibility in both adolescent and adult rats. Neuroscience. 2017;348:324–334.

18. Carbia C, Cadaveira F, Lopez-Caneda E, Caamano-Isorna F, Rodriguez Holguin S, Corral M. Working memory over a six-year period in young binge drinkers. Alcohol. 2017;61:17–23.

19. Salling MC, Skelly MJ, Avegno E, Regan S, Zeric T, Nichols E, Harrison NL. Alcohol consumption during adolescence in a mouse model of binge drinking alters the intrinsic excitability and function of the prefrontal cortex through a reduction in the hyperpolarization-activated cation current. J Neurosci. 2018;38:6207–6222.

20. Ehlers CL, Liu W, Wills DN, Crews FT. Periadolescent ethanol vapor exposure persistently reduces measures of hippocampal neurogenesis that are associated with behavioral outcomes in adulthood. Neuroscience. 2013;244:1–15.

21. Badanich KA, Fakih ME, Gurina TS, Roy EK, Hoffman JL, Uruena-Agnes AR, Kirstein CL. Reversal learning and experimenter-administered chronic intermittent ethanol exposure in male rats. Psychopharmacology. 2016;233:3615–3626.

22. Dalley JW, Cardinal RN, Robbins TW. Prefrontal executive and cognitive functions in rodents: Neural and neurochemical substrates. Neurosci Biobehav Rev. 2004;28:771–784.

23. Crews FT, Vetreno RP, Broadwater MA, Robinson DL. Adolescent alcohol exposure persistently impacts adult neurobiology and behavior. Pharmacol Rev. 2016;68:1074–1109.

24. Boutros N, Semenova S, Liu W, Crews FT, Markou A. Adolescent intermittent ethanol exposure is associated with increased risky choice and decreased dopaminergic and cholinergic neuron markers in adult rats. Int J Neuropsychopharmacol. 2015;18:1–9.

25. Fernandez GM, Savage LM. Adolescent binge ethanol exposure alters specific forebrain cholinergic cell populations and leads to selective functional deficits in the prefrontal cortex. Neuroscience. 2017;361:129–143.

26. Vetreno RP, Broadwater M, Liu W, Spear LP, Crews FT. Adolescent, but not adult, binge ethanol exposure leads to persistent global reductions of choline acetyltransferase expressing neurons in brain. PLoS One. 2014;9:e113421.

27. Vetreno RP, Bohnsack JP, Kusumo H, Liu W, Pandey SC, Crews FT. Neuroimmune and epigenetic involvement in adolescent binge ethanol-induced loss of basal forebrain cholinergic neurons: Restoration with voluntary exercise. Addict Biol. 2020;25:e12731.

28. Woolf NJ, Hernit MC, Butcher LL. Cholinergic and non-cholinergic projections from the rat basal forebrain revealed by combined choline acetyltransferase and Phaseolus vulgaris leucoaglutinin immunohistochemistry. Neurosci Lett. 1986;66:281–286.

29. Zaborszky L, Csordas A, Mosca K, Kim J, Gielow MR, Vadasz C, et al. Neurons in the basal forebrain project to the cortex in a complex topographic organization that reflects corticocortical connectivity patterns: An experimental study based on retrograde tracing and 3D reconstruction. Cereb Cortex. 2015;25:118–137.

30. Picciotto MR, Higley MJ, Mineur YS. Acetylcholine as a neuromodulator: Cholinergic signaling shapes nervous system function and behavior. Neuron. 2012;76:116–129.

31. Howe WM, Berry AS, Francois J, Gilmour G, Carp JM, Tricklebank M, Lustig C, Sarter MJ. Prefrontal cholinergic mechanisms instigating shifts from monitoring for cues to cue-guided performance: Converging electrochemical and fMRI evidence from rats and humans. J Neurosci. 2013;33:8742–852.

32. Parikh V, Kozak R, Martinez V, Sarter M. Prefrontal acetylcholine release controls cue detection on multiple timescales. Neuron. 2007;56:141–154.

33. Runfeldt MJ, Sadovsky AJ, MacLean JN. Acetylcholine functionally reorganizes neocortical microcircuits. J Neurophysiol. 2014;112:1205–1216.

34. Sarter M, Lustig C, Howe WM, Gritton H, Berry, AS. Deterministic functions of cortical acetylcholine. Eur J Neurosci. 2014;39:1912–1920.

35. Savage LM, Chang Q, Gold PE. Diencephalic damage decreases hippocampal acetylcholine release during spontaneous alternation testing. Learn Mem. 2003;10:242–246.

36. Paul CA, Beltz B, Berger-Sweeney J. The nissl stain: A stain for cell bodies in brain sections. CSH Protoc. 2008;pdb.prot4805.

37. Paxinos G, Watson G. The rat brain in stereotaxic coordinates. Cambridge, MA: Academic Press; 2013.

38. Risher ML, Fleming RL, Risher WC, et al. Adolescent intermittent alcohol exposure: Persistence of structural and functional hippocampal abnormalities into adulthood. Alcohol Clin Exp Res. 2015;39(6):989–997.

39. Spear LP. Adolescent alcohol exposure: Are there separable vulnerable periods within adolescence? Physiol Behav. 2015;148:122–130.

40. Nguyen-Louie TT, Tracas A, Squeglia LM, Matt, GE, Eberson-Shumate S, Tapert SF. Learning and memory in adolescent moderate, binge, and extreme-binge drinkers. Alcohol Clin Exp Res. 2016;40:1895–1904.

41. Prado VF, Roy A, Kolisnyk B, Gros R, Prado MA. Regulation of cholinergic activity by the vesicular acetylcholine transporter. Biochem J. 2013;450:265–274.

42. Kolisnyk B, Al-Onaizi MA, Hirata PH, Guzman MS, Nikolova S, Barbash S, Soreq H, Bartha R, Prado MA, Prado VF. Forebrain deletion of the vesicular acetylcholine transporter results in deficits in executive function, metabolic, and RNA splicing abnormalities in the prefrontal cortex. J Neurosci. 2013;33:14908–20.

43. Martyn AC, De Jaeger X, Magalhães AC, Kesarwani R, Gonçalves DF, Raulic S, Guzman MS, Jackson MF, Izquierdo I, Macdonald JF, Prado MA, Prado VF. Elimination of the vesicular acetylcholine transporter in the forebrain causes hyperactivity and deficits in spatial memory and long-term potentiation. Proc Natl Acad Sci. U.S.A. 2021;109:17651–17656.

44. Kyzar EJ, Zhang H, Pandey SC. Adolescent alcohol exposure epigenetically suppresses amygdala arc enhancer RNA expression to confer adult anxiety susceptibility. Biol Psychiatry. 2019;85:904–914.

45. Varlinskaya EI, Kim EU, Spear LP. Chronic intermittent ethanol exposure during adolescence: Effects on stress-induced social alterations and social drinking in adulthood. Brain Res. 2017;1654(Pt B):145–156.

46. Izquierdo A, Brigman JL, Radke AK, Rudebeck PH, Holmes A. The neural basis of reversal learning: An updated perspective. Neuroscience. 2017;345:12–26.

47. Wilson RC, Takahashi YK, Schoenbaum G, Niv Y. Orbitofrontal cortex as a cognitive map of task space. Neuron. 2014;81:267–279.

48. Galaj E, Kipp BT, Floresco SB, Savage LM. Persistent alterations of accumbal cholinergic interneurons and cognitive dysfunction after adolescent intermittent ethanol exposure. Neuroscience. 2019;404:153–164.

49. Lanoue V, Copper HM. Branching mechanisms shaping dendrite architecture. Dev Bio. 2019;451:16–25.

50. Ballesteros-Yáñez I, Benavides-Piccione R, Bourgeois JP, Changeux JP, DeFelipe J. Alterations of cortical pyramidal neurons in mice lacking high-affinity nicotinic receptors. Proc Natl Acad Sci. U.S.A. 2010;107:11567–11572.

51. Kang L, Tian MK, Bailry CDC, Lambe EK. Dendritic spine density of prefrontal layer 6 pyramidal neurons in relation to apical dendrite sculpting by nicotinic acetylcholine receptors. Front Cell Neurosci. 2015; 9:398.

52. Mychasiuk R., Muhammad A., Gibb R., Kolb B. Long-term alterations to dendritic morphology and spine density associated with prenatal exposure to nicotine. Brain Res. 2013;1499:53–60.

53. Spear LP. Timing eclipses amount: The critical importance of intermittency in alcohol exposure effects. Alcohol Clin Exp Res. 2020;44:806–813.

54. Jury NJ, Pollack GA, Ward MJ, Bezek JL, Ng AJ, Pinard CR, Bergstrom HC, Holmes A. Chronic Ethanol During Adolescence Impacts Corticolimbic Dendritic Spines and Behavior. Alcohol Clin Exp Res. 2017;41(7):1298–1308.

55. Kim A, Zamora-Martinez ER, Edwards S, Mandyam CD. Structural reorganization of pyramidal neurons in the medial prefrontal cortex of alcohol dependent rats is associated with altered glial plasticity. Brain Struct Funct. 2015;220:1705–1720.

56. Trantham-Davidson H, Centanni SW, Garr SC, New NN, Mulholland PJ, Gass JT, Glover EJ, Floresco SB, Crews FT, Krishnan HR, Pandey SC, Chandler LJ. Binge-like alcohol exposure during adolescence disrupts dopaminergic neurotransmission in the adult prelimbic cortex. Neuropsychopharmacology. 2017;42:1024–1036.

57. McGuier NS, Padula AE, Lopez MF, Woodward JJ, Mulholland PJ. Withdrawal from chronic intermittent alcohol exposure increases dendritic spine density in the lateral orbitofrontal cortex of mice. Alcohol. 2015;49(1):21–27.

58. Holmes A, Fitzgerald PJ, MacPherson KP, DeBrouse L, Colacicco G, Flynn SM, Masneuf S, Pleil KE, Li C, Marcinkiewcz CA, Kash TL, Gunduz-Cinar O, Camp M. Chronic alcohol remodels prefrontal neurons and disrupts NMDAR-mediated fear extinction encoding. Nat Neurosci. 2012;15:1359–1361.

59. Lees B, Meredith LR, Kirkland AE, Bryant BE, Squeglia LM. Effect of alcohol use on the adolescent brain and behavior. Pharmacol Biochem Behav. 2020;192:172906.

